# Serial Immunohistochemistry for High-Dimensional Single-Cell Spatial Analysis of Human Kidney Biopsies

**DOI:** 10.64898/2026.08.06.743188

**Authors:** Xiaoping Yang, M. Caleb Marlin, Alessandra Ida Celia, Chen-Yu Lee, Alexandre Cammarata−Mouchtouris, Tayte Stephens, Majood Haddad, Liz Bradshaw, Daksh Saksena, Jill Buyon, Peter M. Izmirly, Chaim Putterman, Diane Kamen, Michelle Petri, Accelerating Medicines Partnership: RA/SLE Network, Judith A. James, Joel M. Guthridge, Andrea Fava, Avi Z. Rosenberg

## Abstract

**Background:** Traditional immunohistochemistry (IHC) with chromogen detection has limited multiplex capacity, detecting at most 4 protein markers per tissue section simultaneously, thereby restricting comprehensive spatial analysis of valuable human biopsies. We developed and validated a robust serial IHC (sIHC) staining method to detect multiple antigens on a single kidney biopsy slide, maximizing data yield for diagnosing and studying complex kidney diseases.

**Methods:** Formalin-fixed, paraffin-embedded kidney biopsy sections were subjected to repeated IHC/imaging cycles with antibody removal using an optimized sodium dodecyl sulfate-glycerol buffer stripping protocol. Images were then co-registered, and analysis was performed using a variety of methodologies, including color deconvolution, cell segmentation, and spatial clustering.

**Results:** This optimized sIHC method successfully detected up to 20 antigens on a single slide. Combining image analysis and artificial intelligence software, for example with HALO (Indica Labs), the assay assembles high-dimensional images and enables quantitative histology and single-cell spatial analysis. Using this advanced method, we were able to identify rare cell populations, such as double-negative T cells, that are challenging to detect conventionally.

**Conclusion:** We have developed a validated, high-capacity sIHC protocol that uses standard IHC procedures with commercially available, clinically validated off-the-shelf antibodies. This method is a valuable, cost-effective tool for obtaining extensive, high-dimensional single-cell-resolved spatial data from limited pathology samples, such as a human kidney biopsy.

## Introduction

Immunohistochemistry (IHC) is a foundational technique in the histochemical clinical and experimental assessment of tissues, leveraging antibody-antigen binding detected with a chromogenic reaction to detect specific protein targets within preserved tissue architecture. This approach is practical and cost-effective, making it pivotal for routine diagnostic, prognostic and theragnostic applications. A major limitation of traditional IHC, however, is its capacity, for the most part in routine applications, to detect only one marker per tissue section. This constraint significantly limits the information that can be obtained from limited patient samples, for example, when a comprehensive analysis of the complex immune microenvironment is required. To overcome this, several multiplexing techniques have emerged, including multiplexed Immunohistochemistry/Immunofluorescence (mIHC/mIF)^1,2^, detection of multiple antigens with different chromogens^1,3^, multiplexed ion beam imaging (MIBI)^4^, and imaging mass cytometry (IMC)^5^. While these methods allow simultaneous detection of multiple markers, they often involve trade-offs, including high cost, complex instrumentation, and a limited panel size. Furthermore, many of these platforms require extensive and rigid panel optimization; the addition or replacement of even a single marker often necessitates a re-validation of the entire antibody set, a process that is both time-consuming and resource-intensive. These factors frequently render them cost-prohibitive and inflexible for routine clinical or large-scale research laboratories.

Serial immunohistochemistry (sIHC), also known as multiplexed immunohistochemistry consecutive staining on a single slide (MICSSS)^6^, is a powerful and accessible approach for high-dimensional tissue analysis. This technique employs sequential cycles of antibody staining and stripping on a single tissue section to generate multiplexed IHC images. We present a refined sIHC protocol optimized for routine pathology laboratories that uses reagents and workflows similar to standard IHC and can be performed using either manual or automated platforms. A key innovation of our approach is the use of a gentle stripping buffer, rather than harsh antigen retrieval solutions^7^, to effectively remove prior antibodies while minimizing tissue damage and loss across repeated cycles. Coupled with HALO image analysis (Indica Labs) enabling single-cell level analysis in structurally complex tissue, this methodology supports robust detection of more than 20 antigens with diverse subcellular localization (nuclear, cytoplasmic, and membrane) on a single slide. The resulting high-dimensional images enable quantitative histology, identification of immune cell subtypes, and spatial resolution of the immune microenvironment, facilitating comprehensive single-cell and spatial analysis of immune cells in human kidney biopsy samples.

The utility of this refined sIHC protocol was further validated through its implementation within the Accelerating Medicines Partnership Rheumatoid Arthritis and Systemic Lupus Erythematosus (AMP RA/SLE) network. By applying this platform to a cohort of human kidney biopsies, we successfully generated high-dimensional spatial data across nearly two million individual cells. This large-scale application underscores the capacity of the method for robust immune phenotyping and automated tissue classification, demonstrating its value for sophisticated spatial analysis in the context of complex autoimmune diseases where clinical specimens are often limited and precious.

## Materials and Methods

### Tissue Samples

Human tonsil and control kidney samples were sourced from the Johns Hopkins Pathology tissue bank. Formalin-fixed paraffin-embedded (FFPE) renal biopsy tissue blocks were obtained from a representative subset of Phase 2 lupus nephritis patients enrolled across participating Accelerating Medicines Partnership (AMP) RA/SLE network clinical sites, including Johns Hopkins University (JHU), New York University (NYU), Medical University of South Carolina (MUSC), and Albert Einstein College of Medicine, as previously described^8^ under protocols approved by Institutional Review Boards (IRB). Tissues and biopsy samples were sectioned at a nominal thickness of 5 μm (range 4–6 μm) to ensure structural integrity throughout the iterative staining and stripping cycles.

### Reagents

Primary antibodies and panel specifications are detailed in Supplemental Table 1 (Panel 1) and Supplemental Table 2 (Panel 2). Panel 1 was deployed for primary multi-marker spatial validation (Figure 4), whereas Panel 2 was used for expanded phenotyping of infiltrating immune populations (Figure 5). Specific reagents, including ImmunoDNA retrieve citrate solution (BSB 0022), ImmunoDNA retrieve EDTA solution (0032), peroxidase blocker (BSB 0054), antibody diluent (BSB 0041), ImmunoDNA washer (BSB 0150), and Polydetector Plus Link and HRP (BSB 0270), were purchased from Bio SB Inc. (Santa Barbara, CA). The HRP-conjugated Goat IgG antibody (VC-004-025) was sourced from R&D System (Minneapolis, MN). Vecta Mount AQ mounting medium (H-5501) and the ImmPACT AEC Substrate kit (SK-4205) were purchased from Vector Laboratories (Burlingame, CA). The Periodic Acid Schiff kit and Hematoxylin (Richard Allan Scientific, San Diego, CA) were obtained from VWR Inc. (84000-252).

Stripping Buffer Stocks:

- 10% Sodium dodecyl sulfate (SDS) was prepared in the laboratory.
- 0.5M Tris-HCI (pH 6.8) was prepared in the laboratory.
- â-mercaptoethanol (β-ME) (63689) was purchased from Sigma-Aldrich, St. Louis MO.
- Glycerol solution (G9012) was purchased from Sigma-Aldrich.

Equipment and analysis software:

- Lab oven - Quincy lab Inc. (Fisher Scientific, S50170). The oven was placed within a chemical hood in the laboratory.
- Pressure cooker - Cuisinart pressure cooker (any commercial vendor)
- MoticEasyScan Pro scanner (Motic, Vancouver, BC, Canada)
- Slide baker (Thermo scientific, Waltham, MA)
- HALO (Indica Labs, Albuquerque, NM)

### Serial Immunohistochemistry (sIHC) Protocol

#### Day 1: Deparaffinization, Antigen Retrieval, and IHC

Slides were baked at 60°C for 1 h, deparaffinized in xylene (3 × 8 min), and rehydrated through graded ethanol (100%, 90%, 70%; 2 min each), followed by rinsing in distilled water (dH₂O). Antigen retrieval was performed in 1× citrate buffer (pH 6.0; 200 mL) using a pressure cooker at high pressure for 18 min. Slides were cooled for 15 min at room temperature and rinsed in 4-5 changes of dH₂O to ensure the complete removal of residual detergent (Tween 20).

For IHC, slides were equilibrated in wash buffer (Bio SB ImmunoDNA Washer) for 1 min, incubated with peroxidase blocker (Bio SB) for 5 min, and washed once in dH₂O and once in wash buffer (1 min each). Primary antibodies diluted in antibody diluent (Bio SB) were applied for 90 min at room temperature, followed by one rinse in dH₂O and two washes in wash buffer (3 min each). Poly-Detector Plus Link (Bio SB, BSB-0270) or a species-appropriate HRP-conjugated secondary antibody was applied for 15-20 min, followed by one dH₂O rinse and two wash buffer washes (3 min each). Poly-Detector Plus HRP (Bio SB, BSB-027) was then applied for 15 min, followed by identical washes.

Chromogenic detection was performed using AEC (Vector Laboratories) for 10-25 min, followed by one wash in wash buffer and two washes in dH₂O (3 min each). Slides were counterstained with hematoxylin (Richard Allan) for 3 min, washed in tap water for 3 min, mounted with aqueous mounting medium (VectaMount AQ), coverslipped, and dried for 1 h at room temperature or overnight at 4°C. Slides were scanned (MoticEasyScan) and stored at 4°C.

#### Day 2: Coverslip Removal, Antibody Stripping, and Restaining

Coverslips were removed by incubating slides in dH₂O at 50°C until detachment (∼20 min). Slides were rinsed in dH₂O, incubated in 70% ethanol for 2 min, 90% ethanol for 15 min, returned to 70% ethanol for 2 min, and rinsed in dH₂O.

Antibody stripping was performed using stripping buffer (2% SDS, 62.5 mM Tris-HCl pH 6.8, 0.8% β-mercaptoethanol, 5% glycerol). Slides were incubated in stripping buffer at 60°C for 20 min. Alternatively, stripping was performed at 55°C for 30 min in a water bath or by steaming for 10 min. Slides were rinsed three times in dH₂O (10 min each), washed twice in wash buffer (10 min each), and equilibrated in PBS for 10 min. Subsequent IHC staining was performed as described for Day 1.

#### Day 3 and Subsequent Cycles

Day 2 procedures were repeated for each additional staining cycle. A schematic overview of the sIHC workflow is shown in Figure 1A.

**Figure 1.**
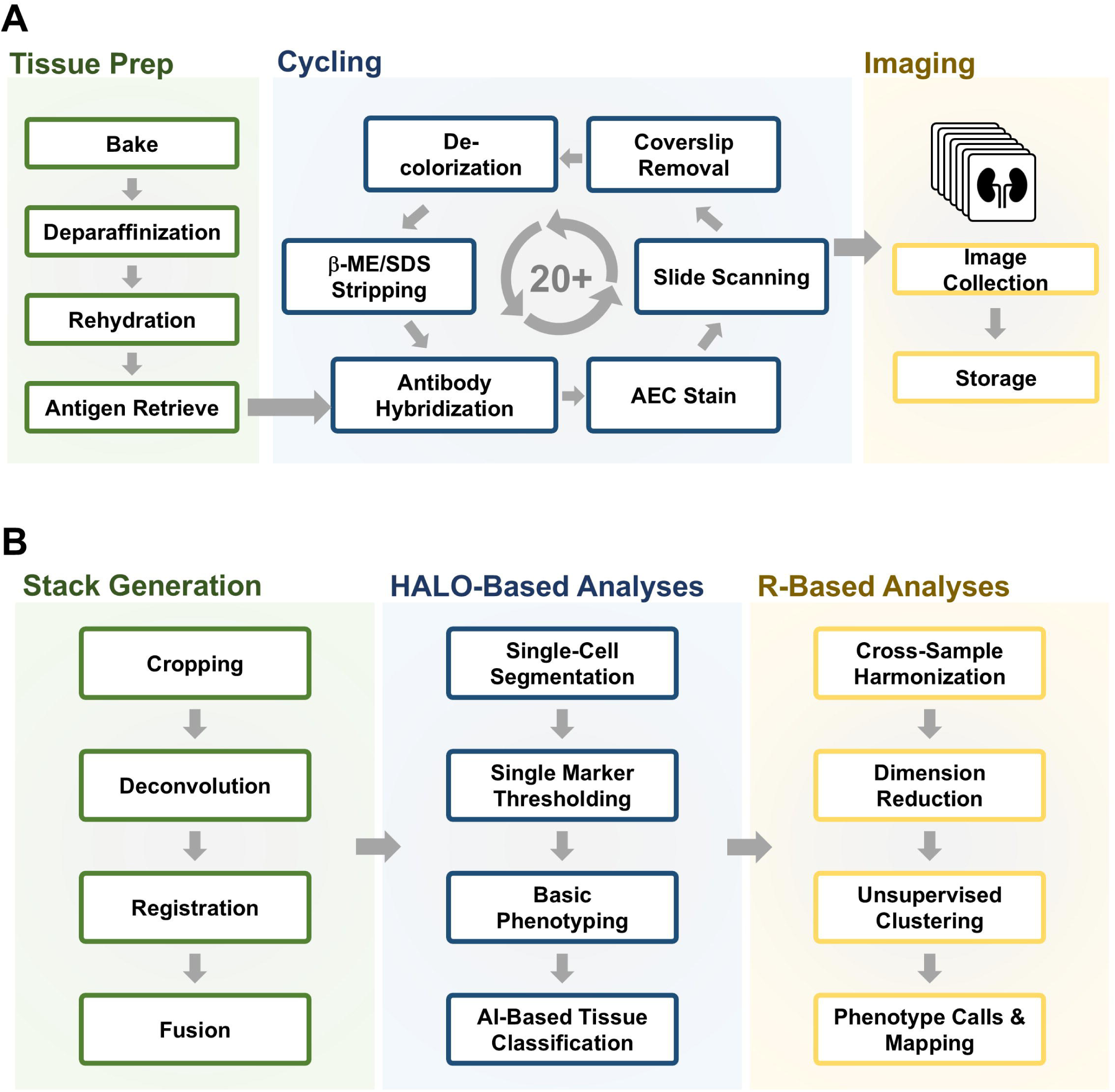
sIHC Protocol and Analysis Workflow Overview. (A) sIHC Workflow. On the first day, staining is performed using the standard immunohistochemistry procedure. From the second day onwards, the slide undergoes repeated cycles that include coverslip removal, de-colorization, antibody stripping, re-hybridization with new antibodies, staining, and image acquisition. All individual images are collected on one slide. (B) Data Analysis Workflow. Raw images undergo Stack Generation (Cropping, Deconvolution, Registration, and Fusion) to produce a single, multi-layer image stack. This stack is then subjected to HALO-Based Analyses (Single-Cell Segmentation, Single Marker Thresholding, Basic Phenotyping, and AI-Based Tissue Classification). Extracted single-cell data proceeds to R-Based Analyses (Cross-Sample Harmonization, Dimension Reduction, Unsupervised Clustering, and Phenotype Calls & Mapping).

### Constructing the Image Stacks, Tissue Classifier, and Image analysis

Image files were imported into HALO-AI v3.5 (Indica Labs) for deconvolution, pseudo-coloring, co-registration, alignment, and fusion into a single composite image containing all markers, a representative DNA channel, and the counterstain. Tissue regions were identified using a DenseNet V2 AI tissue classifier trained on an original co-registered PAS image. Individual cells were detected via nuclear identification and segmented using cell distance boundaries.

Single-cell marker expression data were analyzed in R (version 4.2). Cells located at tissue edges or within regions classified as wrinkles were excluded. Marker expression values were normalized using the centered log-ratio (CLR) transformation and standardized within each sample. Data were harmonized across samples using the Harmony algorithm^9^ to minimize batch effects. Dimensionality reduction was performed using principal component analysis (PCA) followed by uniform manifold approximation and projection (UMAP).

Cell clustering was performed using k-nearest neighbors (KNN) and shared nearest neighbor (SNN) algorithms. Immune cell clusters were identified by CD45 expression levels greater than 2 standard deviations above the mean, while clusters with high keratin expression were excluded to remove renal tubular epithelial cells. The remaining immune populations were re-clustered using KNN and SNN to resolve distinct immune cell types.

Clustered data were visualized using heatmaps and feature plots. Heatmaps incorporated hierarchical clustering with dendrograms applied to both rows and columns to illustrate relationships among samples and marker expression profiles. A summary of the analysis workflow is shown in Figure 1B.

### Identification of Double Negative T-Cells

Marker thresholds were set for CD45, CD3, CD4, and CD8 in the HALO imaging analysis software within the fully stacked and registered images. Using the results tab, the software identifies and highlights cells using basic logic commands based upon the resulting gates. For example, we designed a phenotype profile that contained positive staining for CD45 and CD3, but did not contain positive staining for CD4 and CD8. These cells were identified and highlighted withing the HALO software.

## Results

### Complete Primary Antibody Elution with SDS Stripping Buffer

We validated the efficiency of antibody stripping by comparing a conventional heat-based antigen retrieval solution (citrate buffer)^10^ with our SDS-based stripping buffer^7^ for removal of bound primary antibodies. Tissue sections were first stained with CD20 and then either boiled in citrate retrieval buffer for 15 min or incubated in SDS stripping buffer at 60°C for 20 min. Citrate-based retrieval failed to fully remove the primary antibody, leaving a significant residual CD20 signal. In contrast, the SDS stripping buffer achieved complete antibody elution with no detectable residual staining (Figure 2A). Effective stripping was shown to be essential for accurate subsequent staining cycles. Sections stripped with SDS buffer allowed for clean, unambiguous restaining with CD56. However, incomplete stripping with citrate buffer resulted in mixed, uninterpretable co-localization of CD20 and CD56 signals (Figure 2B). This technical superiority extended to nuclear and shared markers; while EDTA retrieval failed to fully strip CD45, SDS stripping enabled complete removal, allowing for clear FoxP3 staining in the following cycle (Figure 2C).

**Figure 2.**
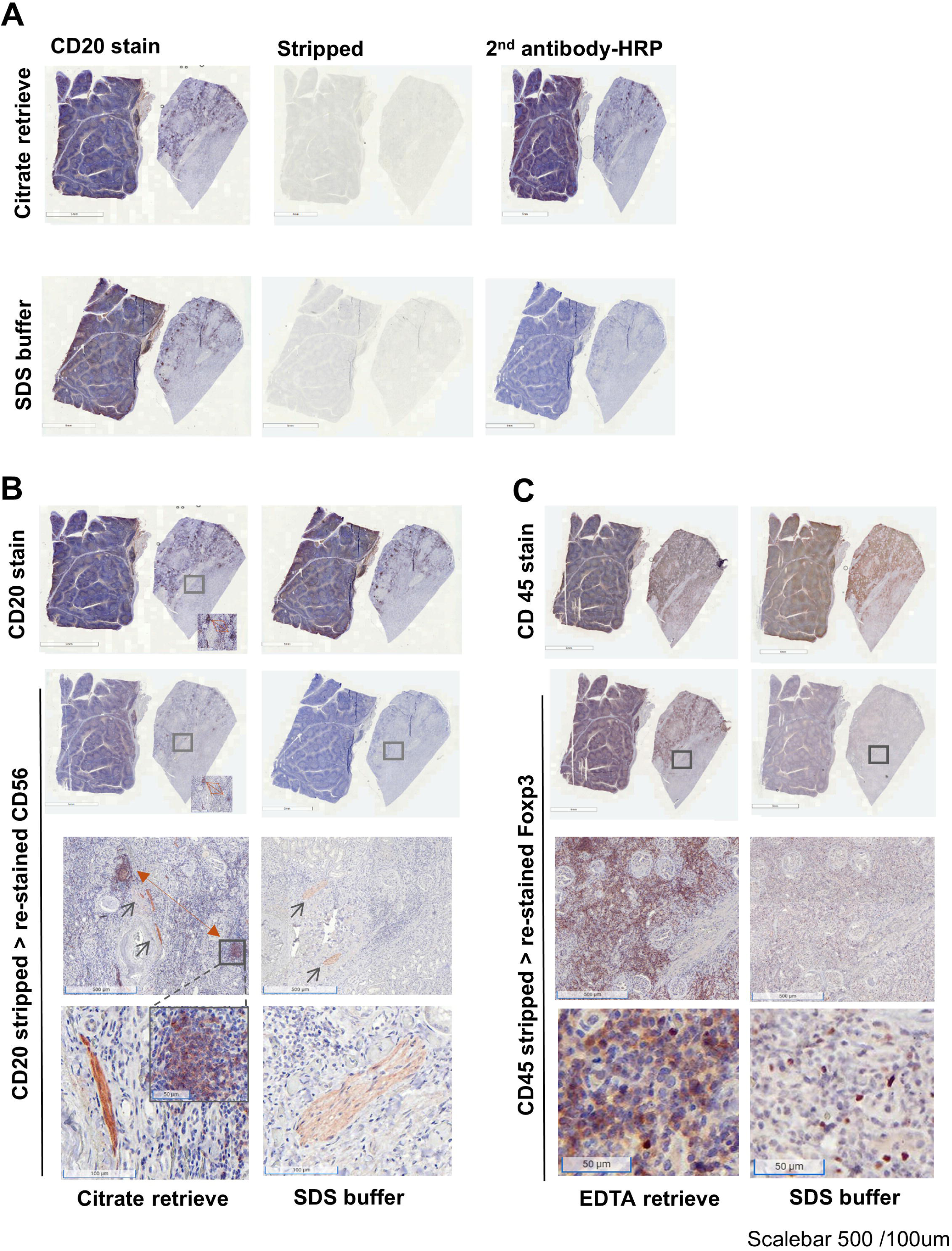
Complete Removal of Antibody with SDS Stripping Buffer. (A) Comparison of Antibody Elution Efficacy. Tonsil (left) and kidney (right) tissue sections (scale bars: 5 mm) were initially stained for CD20 (left panels). Following elution with citrate retrieval solution (middle up) or SDS buffer (middle low), The slides were re-incubated with secondary reagent (HRP) without a primary antibody (right panels). The images show residual staining in the citrate-retrieval treated slide but not in SDS buffer treated slide (right panels). This indicated that the citrate-treated section retained residual primary antibody, while the SDS section did not. (B) SDS Stripping Ensures Accurate Re-staining. After initial CD20 staining (top panels), slides were stripped and re-probed with CD56 (middle and lower panels). The SDS-treated slide (right) shows clean CD56 stains (black arrows), while the citrate-treated slide (left) exhibits mixed stains, with residual CD20 signal (red arrows) co-localizing with the CD56 signal. (C) Efficacy for Nuclear Antigens. Slides initially stained for CD45 (upper panels) were stripped with EDTA or SDS buffer and re-hybridized with FoxP3 (middle and low panels). The EDTA-treated slide shows mixed FoxP3 and residual CD45 stains, while the SDS-treated slide shows sole FoxP3 stains. Scale bars 5mm, 500 µm and 50 µm respectively.

### Mitigation of Tissue Damage and Optimization with Glycerol

While highly effective for elution, repeated stripping cycles carry the risk of cumulative tissue degradation. To enhance protocol robustness, we optimized the stripping buffer by supplementing it with glycerol. Over 18 sequential cycles, samples processed without glycerol exhibited progressive tissue damage and significant loss (Figure 3A, upper panel). Conversely, glycerol-treated samples maintained structural integrity, preserved cellular architecture, and demonstrated superior staining intensity even after 18 cycles (Figure 3A, lower panel).

**Figure 3.**
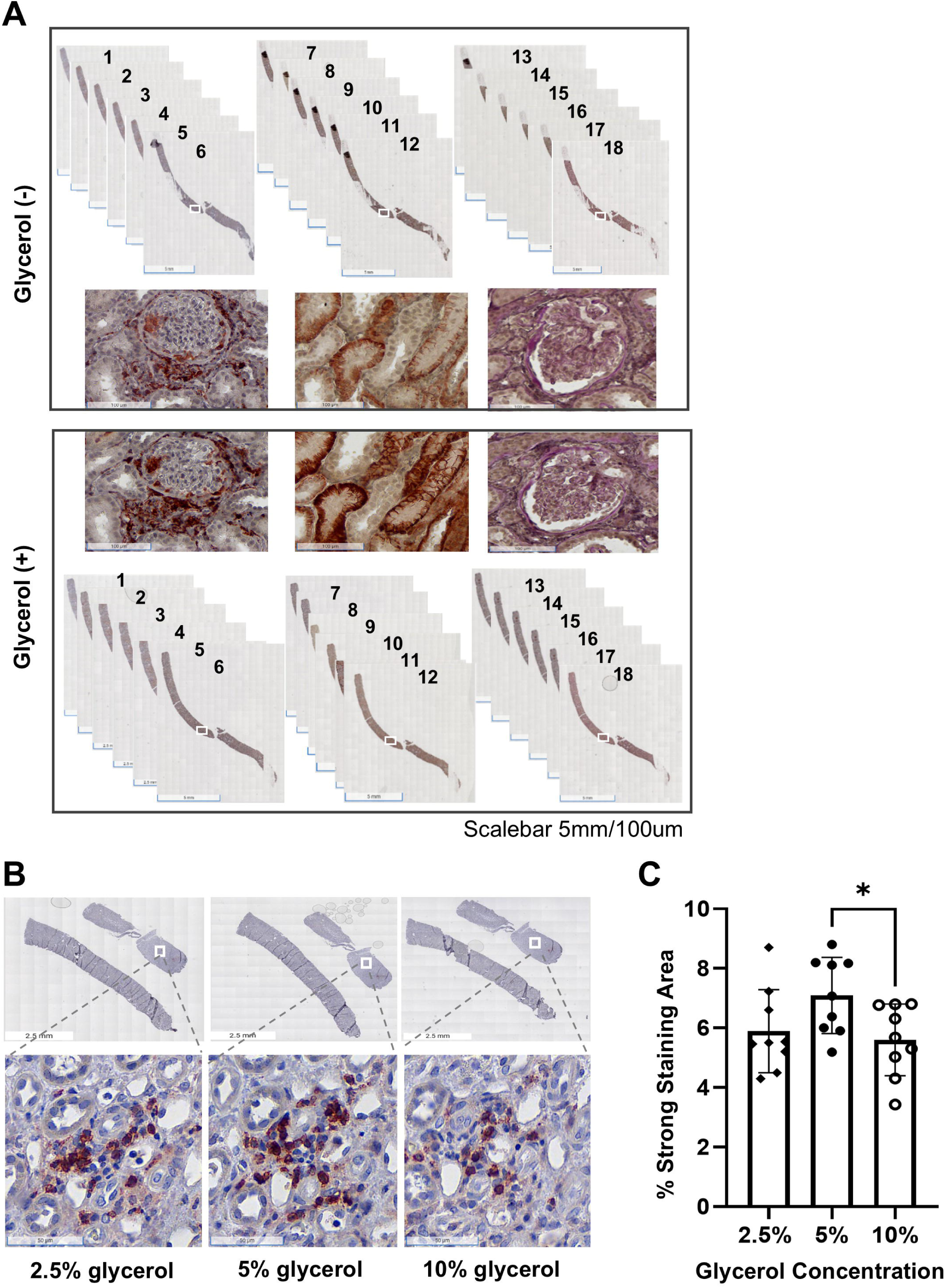
Glycerol Prevents Tissue Loss and Enhances Staining Robustness. (A) Glycerol Mitigates Tissue Damage and Loss. A pair of kidney biopsy slides were subjected to 18 cycles of sIHC. Whole biopsy images (scale bar: 5 mm) and corresponding magnification images (scale bar: 100 μm) are shown over the cycles. The upper panels (Glycerol(−)) show significant tissue damage and loss that begins in early cycles and worsens over time. In contrast, the lower panels (Glycerol(+)) show minimal tissue loss and maintain richer positive staining and clear cellular architecture after 18 cycles. (B) Optimal Glycerol Concentration. Biopsy samples from nine subjects were stripped with SDS buffer containing 2.5%, 5%, or 10% glycerol between CD8 staining cycles. (C) The graph summarizes the percentage of strong staining area quantified by HALO Area-Quantification-v2.4 analysis. The 5% glycerol concentration resulted in the strongest staining, while 10% glycerol showed a significant decrease in staining area compared to 5% (p<0.05, Student’s t-test).

Optimization experiments identified 5% glycerol as the ideal concentration, yielding a significantly greater stained area compared to 2.5% or 10% glycerol (Student’s t-test, p < 0.05; Figure 3B and 3C). This was quantified using HALO area-quantification analysis following sequential CD8 staining and stripping. Supporting long-term antigenicity, robust staining was maintained for the membrane antigen CD138 through 10 cycles and the cytoplasmic antigen PTPRO through 8 cycles (Supplemental Figure 1).

### High-Dimensional Image Construction and sIHC Tissue Architecture Analysis

Using the optimized SDS/glycerol stripping method, we successfully generated high-dimensional images from single kidney biopsy sections using a 20-antigen panel (Panel 1; Supplemental Table 1). This panel encompassed 13 transmembrane markers (such as CD3, CD4, CD8, CD16, CD20, CD45), four cytoplasmic antigens (KIM-1, PTPRO, Pan-Keratin, α-SMA), and three nuclear antigens (Ki-67, cCas3, FoxP3), alongside a PAS counterstain to highlight underlying tissue architecture.

Raw brightfield images were processed using the HALO analysis workflow for deconvolution, registration, and fusion (Figure 4A). The integration of these 20 registered images created a single multi-layer stack that resolved complex renal microenvironments (Figure 4B, i). Specific marker combinations enabled the precise definition of anatomical structures: PTPRO and CD34 delineated the glomerulus, Keratin and KIM-1 defined renal tubules, and α-SMA/Keratin identified fibrotic or lymphoid regions (Figure 4B, ii). Furthermore, the panel allowed for the spatial mapping of infiltrating lymphocytes (CD3, CD4, CD8, FoxP3) and the distribution of macrophage subsets (CD36, CD163, CD169) throughout the tissue (Figure 4B, iii-iv).

**Figure 4.**
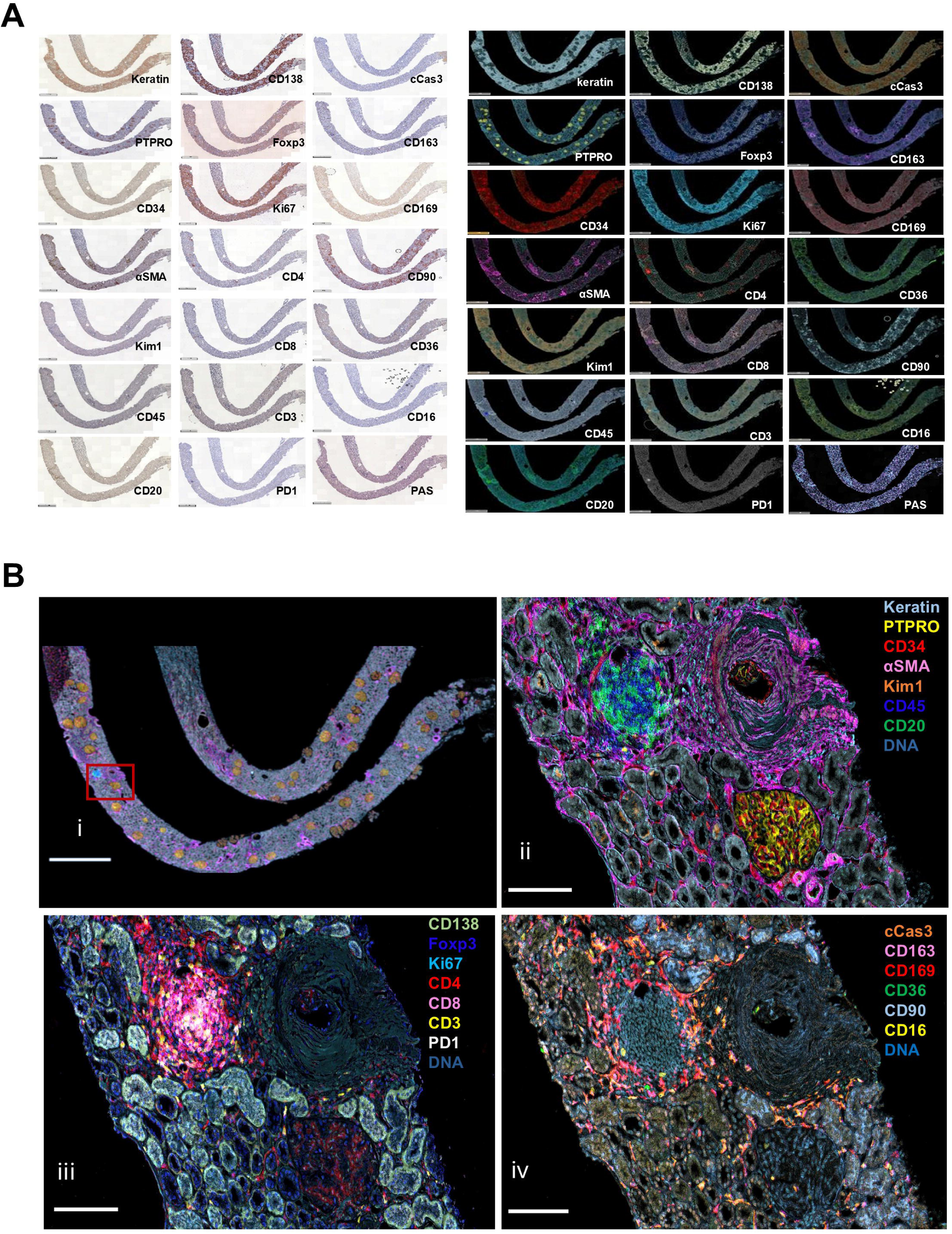
Generation of High-Dimensional Images with Multiple Markers. (A) Image Stack Generation. The sIHC cycles produce a series of raw brightfield images (20 IHC images + PAS). The HALO software is used to deconvolute these raw images into pseudo-color images of the 20-plex panel showing distinct localization of nuclear, cytoplasmic, and membrane-bound antigens. (B) (i) Fused Image and corresponding square for feature mapping and architecture. (ii) Fibrotic Renal Structure and Lymphoid Regions. Specific markers define anatomical structures: α-SMA and keratin indicate scar tissue; PTPRO and CD34 outline the glomerulus; CD45 and CD20 delineate lymphoid regions; and KIM−1 outlines renal tubules. (iii) Lymphocyte Infiltration. Markers such as FoxP3, Ki−67, PD−1, CD3, CD4, and CD8 identify various lymphocytes within the infiltrated lymphoid regions. CD138 outlines the tubular membrane. (iv) Macrophage Distribution. Macrophage markers CD163, CD169, CD36, and CD16 show macrophages distributed around the lymphoid regions. CD90 outlines proximal tubules, and cCas3 markers show apoptotic cells.

### Validation within the AMP RA/SLE Network and AI-Powered Classification

To demonstrate the translational utility of the sIHC platform, we applied the 20-plex panel (Panel 2; Supplemental Table 2) to kidney biopsies within the AMP RA/SLE network (Figure 5A). High-resolution mapping confirmed the maintenance of a high signal-to-noise ratio and precise subcellular localization across large-scale clinical samples (Figure 5B–C).

**Figure 5.**
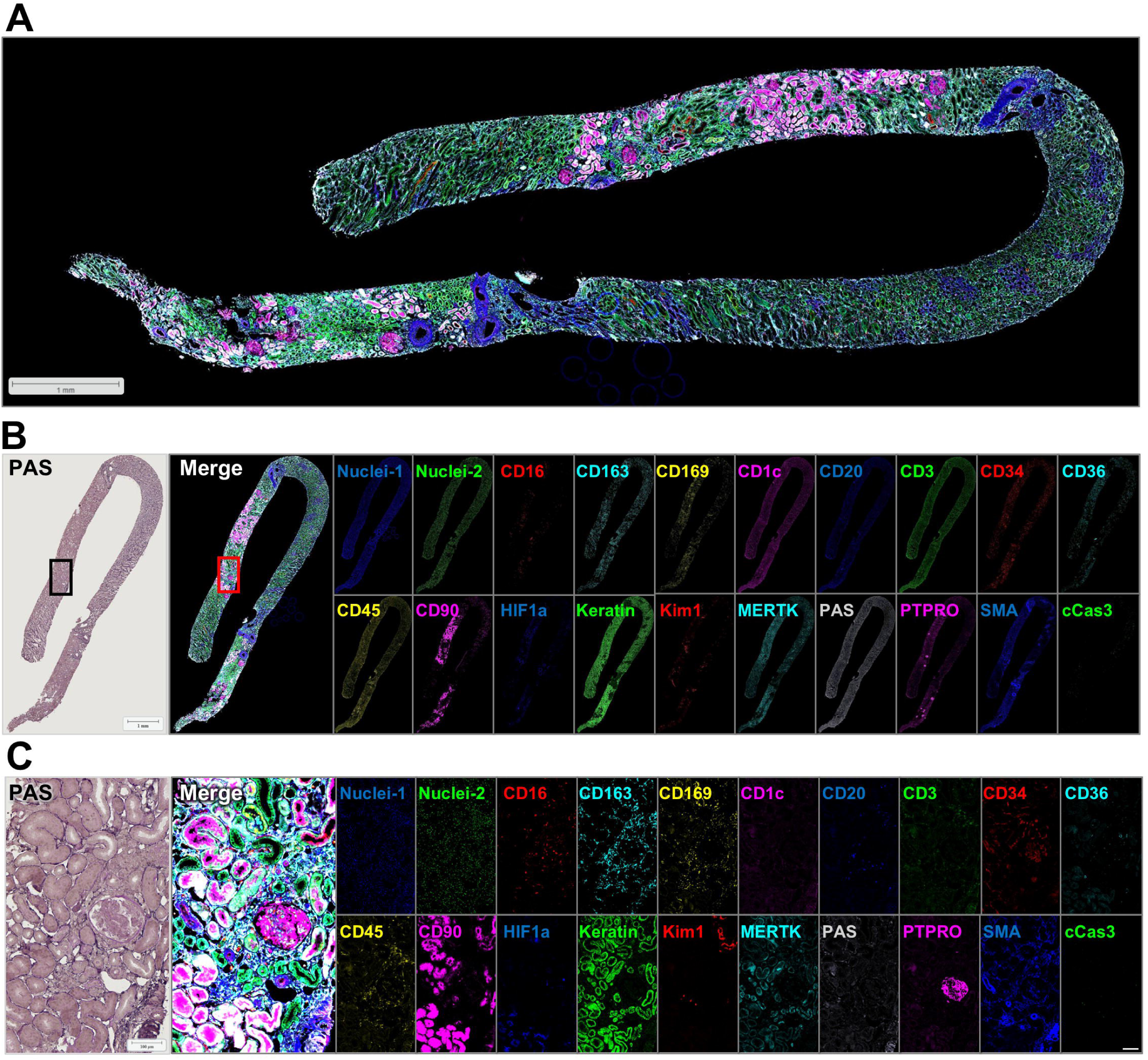
Comprehensive Panel Visualization in Lupus Nephritis. (A) Low-magnification, large view of all channels aligned, registered, and merged on a Lupus Nephritis kidney biopsy from the AMP RA/SLE network. (B) Low-magnification overview of a representative kidney biopsy from the AMP RA/SLE project. The PAS-stained section (left) is shown alongside a multi-marker composite (merge) image (right), illustrating the spatial organization of the immune microenvironment across the whole tissue section. (C) High-magnification detail of the regions indicated in (B). The individual channels and composite images highlight the precise subcellular localization (nuclear, cytoplasmic, and membrane) and the high signal-to-noise ratio maintained throughout the multiplexing cycles. Scale bars: 1 mm and 100 µm respectively.

We used the HALO DenseNet V2 AI tissue classifier to automatically segment specific regions, including glomeruli, interstitial tissue, and proximal/distal tubules (Figure 6A). For broad immune phenotyping, we analyzed 29 human lupus nephritis biopsies profiled with an expanded 36-protein marker set, identifying a total of 1,913,845 individual cells. Of these, 182,783 CD45+ leukocytes were further classified into 10 distinct immune cell populations via unsupervised clustering and UMAP dimension reduction (Figure 6B).

**Figure 6.**
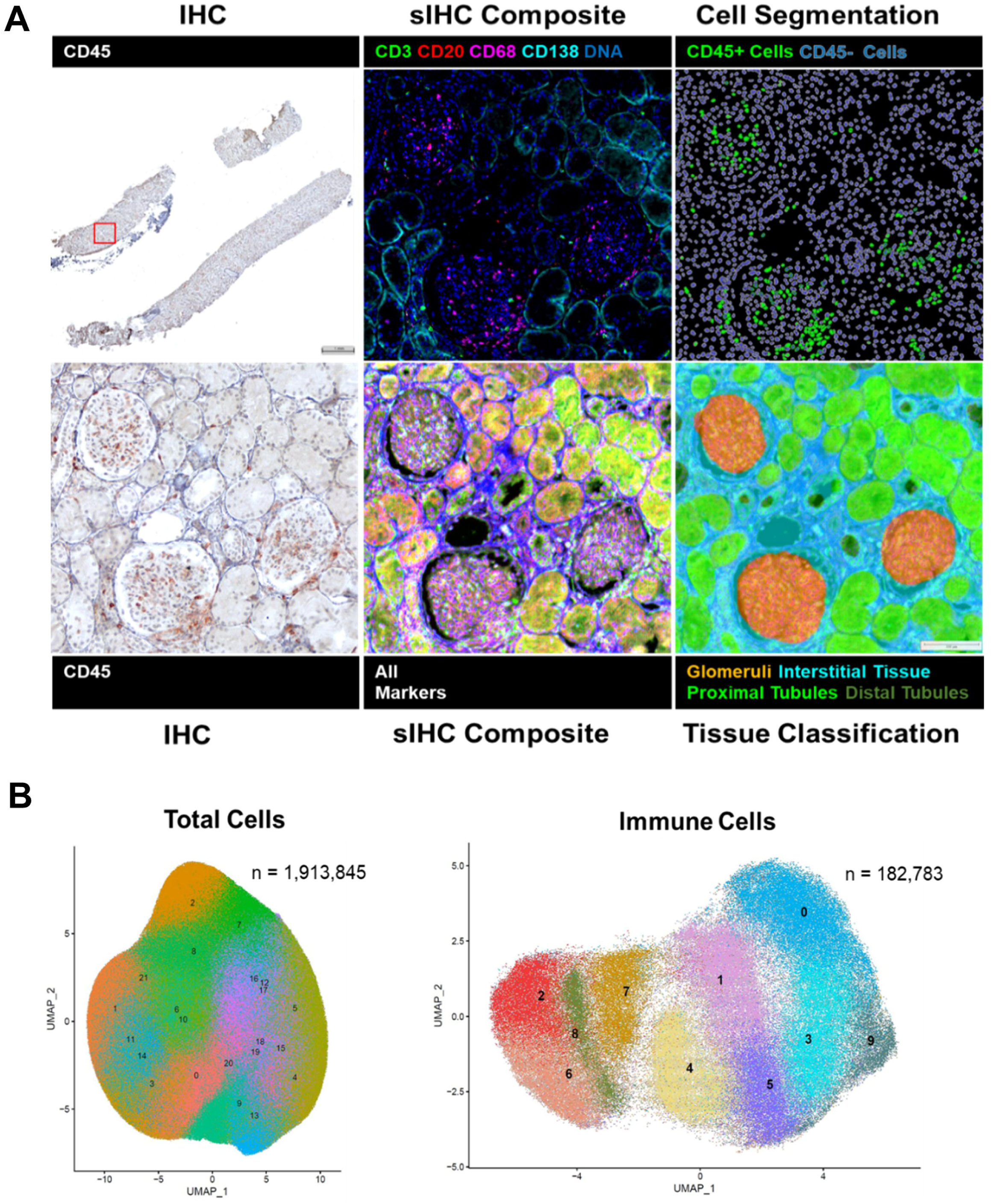
Tissue Classification and Single-Cell Data Analysis. (A) Tissue Classification and Cell Segmentation. The HALO DenseNet V2 AI tissue classifier clearly identifies and segments tissue regions, including glomeruli, proximal tubules, distal tubules, and interstitial tissue. The sIHC composite image shows multiple markers overlaid (CD3, CD20, CD68, CD138, DNA). (B) Single-Cell Spatial Analysis. The analysis included 29 human kidney biopsies with 36 protein markers. A total of 1,913,845 cells were identified. Of these, 182,783 CD45+ cells were classified into 10 distinct immune cell populations via unsupervised clustering and UMAP dimension reduction.

### Spatial Identification of Rare Double-Negative T Cells

We leveraged the platform’s high-dimensional capacity to identify rare and phenotypically challenging cell populations, specifically Double-Negative T cells (DNTs: CD45+, CD3+, CD4-, CD8-). Using HALO thresholding, we successfully distinguished DNTs from Helper (CD4+), Cytotoxic (CD8+), and Double-Positive (CD4+, CD8+) T-cell subsets. The workflow allowed for the spatial localization of individual DNTs within the tissue (Figure 7, upper panel). This phenotypic identification was visually validated by comparing deconvolved pseudo-color images with the original raw brightfield IHC stains. To ensure the accuracy of the Double-Negative (CD3+CD4-CD8-) call, we used a sequential gating strategy within the registered image stack, where thresholds for positivity were defined based on the signal-to-noise ratio of each individual cycle. This allowed us to confirm CD3 positivity in the absence of CD4 and CD8 expressions within the same coordinate-mapped cell boundaries (Figure 7, middle and lower panels). Furthermore, the spatial context of these cells was cross-referenced with the tissue classifier to ensure they were located within expected T-cell niches, rather than representing non-specific staining in damaged tissue.

**Figure 7.**
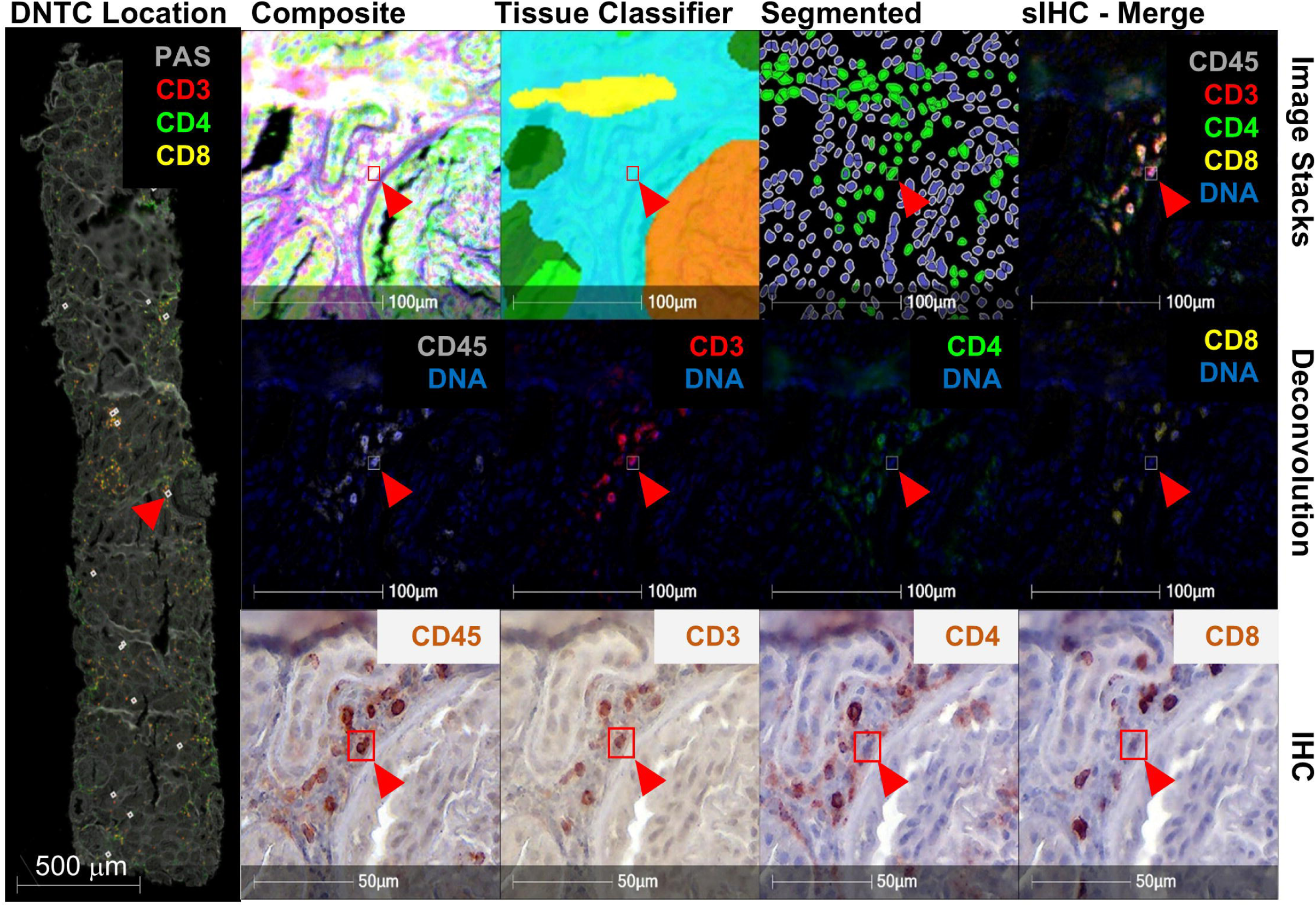
Spatial Localization of Double-Negative T Cells. Identification of Double-Negative T Cells (DNT). DNTs (CD45+, CD3+, CD4-, CD8-) were identified using the phenotype function in HALO v3.5. A representative DNT (red arrowhead) is localized across the composite image, tissue classifier, and segmented image (upper panel). A composite of the deconvolved image stacks (middle panel) and the original unedited IHC staining (lower panel) confirm CD45 and CD3 positivity but lack of CD4 and CD8 expression.

## Discussion

sIHC depends on the complete removal of primary antibodies between cycles to prevent false-positive signals. Existing protocols often rely on antigen retrieval methods for elution, which generally yield clear results over a limited number of cycles (typically 7-8)^10,11^ but frequently fail with high-affinity monoclonal antibodies and the stable cross-linking associated with secondary reagents^7^. Here, we confirmed that conventional retrieval methods resulted in inconsistent and incomplete stripping. To address this issue, Gendusa *et al.*^7^ tested three harsh elution buffers: a pH 2 glycerol buffer, a 6 M urea hot buffer, and a β-ME/SDS buffer. They found that the β-ME/SDS buffer was superior. We adopted this buffer and found that it completely removed bound primary antibodies without affecting the binding of new antibodies to antigens. Although the β-ME/SDS buffer is highly effective, repeated exposure significantly increases the risk of tissue damage and detachment, a major limitation for scarce biopsy samples. Our key technical innovation is the incorporation of 5% glycerol into the stripping buffer. As a protein stabilizer, this addition transformed the methodology, significantly mitigating tissue loss and damage across repeated cycles. Glycerol likely acts as a cytoprotectant by stabilizing protein-protein interactions and reducing the dielectric constant of the stripping solution, thereby preventing the excessive tissue swelling and basement membrane detachment typically associated with hot SDS/β-ME buffers. This optimized protocol enabled up to 20 reliable staining cycles, a substantial increase that allows the collection of extensive, multiplexed information from a single biopsy slide. While tissue detachment and long processing time remain practical limitations, our protocol represents a significant advancement in maximizing data yield from limited clinical samples.

The strategic coupling of our robust sIHC protocol with advanced HALO pathology analysis software yields a high-dimensional dataset, positioning our work as a valuable, accessible alternative to expensive imaging platforms. This combined approach goes beyond simple visualization, enabling the creation of clear, composite images that accurately classify tissue structures and identify cell types based on comprehensive antigen profiles, thereby allowing direct visualization of pathological morphology and the immune microenvironment in a single image. The platform facilitates rigorous quantitative histology by integrating cell segmentation with single-cell marker data, making high-content, single-cell protein analysis accessible to standard pathology laboratories. The power of this platform is demonstrated by how we successfully identified and localized DNTs, an immunologically relevant subset difficult to phenotype using conventional methods^12^.

In conclusion, we have developed an optimized, robust sIHC protocol capable of staining up to 20 protein markers on a single FFPE tissue slide. The incorporation of a 5% glycerol-enhanced stripping buffer is the crucial factor that increases the number of usable cycles while maintaining essential tissue integrity. Coupled with HALO AI analysis, this method enables high-dimensional quantitative histology and single-cell spatial phenotyping of complex tissue microenvironments, which is invaluable for studies relying on small or irreplaceable biopsy samples. Future efforts will focus on automating the staining and stripping steps to reduce overall processing time and enhance throughput for large-scale research and clinical applications.

## Author Contributions

X.P.Y. designed and performed sIHC, collected and analyzed raw data, and wrote the first draft. M.C.M. processed images, analyzed data, and wrote the first draft. A.I.C. performed sIHC, collected and analyzed data, and wrote the first draft. A.C.M. extensively revised and edited the manuscript. T.S., M.H., and L.B. analyzed data. J.B., C.P., D.K., M.P., J.J., and J.G. were a part of the Accelerating Medicines Partnership in RA/SLE that contributed samples and resources to this work. A.F. and A.Z.R. collected biopsy samples, supervised the experimental design, analyzed data, and wrote and edited the manuscript.

## Data Availability

The datasets generated and analyzed during this study are deposited in the Synapse repository (https://sageb.io/Q7bm1A).

## Funding

This work was supported by the Accelerating Medicines Partnership® Rheumatoid Arthritis and Systemic Lupus Erythematosus (AMP® RA/SLE) Program. AMP is a public-private partnership (AbbVie Inc., Arthritis Foundation, Bristol Myers Squibb, GlaxoSmithKline, Lupus Foundation of America, Lupus Research Alliance, Janssen, Merck Sharp & Dohme Corp., NIH National Institute of Allergy and Infectious Diseases, NIH National Institute of Arthritis and Musculoskeletal and Skin Diseases, Pfizer Inc., Rheumatology Research Foundation, Sanofi, and Takeda Pharmaceuticals International Inc.) created to develop new ways of identifying and validating promising biological targets for diagnostics and drug development. This work was supported by NIH grants UH2-AR067676, UH2-AR067677, UH2-AR067679, UH2-AR067681, UH2-AR067685, UH2-AR067688, UH2-AR067689, UH2-AR067690, UH2-AR067691, UH2-AR067694, UM2-AR067678, and AR074096. The Oklahoma Rheumatic Disease Research Cores Center is funded by NIH P30AR073750. The Hopkins Lupus Cohort is funded by NIH R01 AR069572. AF is supported by NIH R01 DK134625, the Jerome L. Greene Foundation, the Plank Scholarship, and the Lupus Foundation of America G-2104-01274.

## Accelerating Medicines Partnership (AMP) RA/SLE Network Members

Jennifer H. Anolik, William Apruzzese, Arnon Arazi, Yemil Atisha-Fregoso, Jennifer L. Barnas, H. Michael Belmont, Celine C. Berthier, Nicole Bornkamp, Michael B. Brenner, Robert M. Clancy, Carla M. Cuda, Michelle Curtis, Maria Dall’Era, Anne Davidson, Evan Der, Betty Diamond, Eugene Drokhlyansky, Thomas M. Eisenhaure, Derek Fine, Richard A. Furie, Beatrice Goilav, Jennifer Grossman, Hasret Gunduz, Siddarth Gurajala, Nir Hacohen, David A. Hildeman, Jeffrey B. Hodgin, V. Michael Holers, Paul J. Hoover, Alice Horisberger, Mariko Ishimori, Nicole Jordan, Kenneth C. Kalunian, Matthias Kretzler, Blue Lake, Chun-Hao Lee, Joseph Mears, Maureen A. McMahon, Rajasree Menon, Manny Monroy-Trujillo, Fernanda Payan-Schober, Harris Perlman, Michael Peters, James Pullman, Deepak A. Rao, Soumya Raychaudhuri, Raktima Raychowdhury, Brad Rovin, Saori Sakaue, Amit Saxena, Daniel Schwartz, Jennifer Seifert, Ummara Shah, Kamil Slowikowski, Dawn Smilek, Nicholas W. Sugiarto, Patti Tosta, Thomas Tuschl, Paul J. Utz, Michael Weisman, David Wofsy, E. Steve Woodle, Ming Wu, Qian Xiao, and Yu Zhao.

## Declaration of Competing Interest

All authors declare no competing interests.

## Ethics Approval and Consent to Human Samples

The Institutional Review Board (IRB) of Johns Hopkins University approved human autopsy and biopsy samples used for the study.

## Notes

https://sageb.io/Q7bm1A

### Competing Interest Statement

The authors have declared no competing interest.

