## Supplemental Information for "Serial Immunohistochemistry for High-Dimensional Single-Cell Spatial Analysis of Human Kidney Biopsies"

**Supplemental Figure 1. Robust Antigenicity Through Repeated Stripping Cycles.** The figure displays images of two individual slides stained through 10 rounds of stripping and staining cycles. PTPRO, a cytoplasmic antigen (left panels), maintained its staining area through 8 rounds, with a notable reduction only after round 9 and minimal staining by round 10. CD138, a renal tubular membrane antigen (right panels), did not show an obvious reduction in positive staining after 10 rounds; the slight increase in signal intensity observed in later cycles is attributed to cumulative background deposition inherent to the iterative chromogenic process and does not indicate a loss of specificity. Using HALO Area-Quantification-v2.4 analysis, the accompanying table confirms the minimal change in PTPRO positive area up to round 8 and CD138 positive area up to round 10.

**Supplemental Table 1. Antibody Panel 1 Specifications.** Primary antibodies, clone information, dilutions, and targets for the 20-marker panel deployed for multi-marker spatial validation (Figure 4).

| Antibody | Vendor (City, State/Country) | Cat# | Note | Dilution factor | Location |
|---|---|---|---|---|---|
| CD3 | Bio SB (Santa Barbara, CA) | BSB 6452 | RmAb | 1:100 | T-cell |
| CD4 | Bio SB (Santa Barbara, CA) | BSB 5151 | RmAb | 1:100 | T-help cell |
| CD8 | Bio SB (Santa Barbara, CA) | BSB 2848 | RmAb | 1:100 | T-killer cell |
| CD16 | Bio SB (Santa Barbara, CA) | BSB 3324 | RmAb | 1:100 | NK, Neutrophils, MΦ, Glomeruli |
| CD20 | Bio SB (Santa Barbara, CA) | BSB 5193 | MmAb | 1:200 | B-cell |
| CD34 | Bio SB (Santa Barbara, CA) | BSB 6488 | RmAb | 1:100 | Vascular Endothelial |
| CD36 | Thermo Fisher (Waltham, MA) | 18836-AP | RmAb | 1:1000 | MΦ, monocyte, vascular endothelium |
| CD45 | Dako (Carpinteria, CA) | M0701 | MmAb | 1:200 | Tyrosine Phosphatase, all WBC |
| CD90 | Abcam (Cambridge, UK) | ab133350 | RmAb | 1:50 | Proximal tubular & mesenchymal |
| CD138 | Bio SB (Santa Barbara, CA) | BSB 6530 | RmAb | 1:100 | Kidney tubular cells |
| CD163 | Bio SB (Santa Barbara, CA) | BSB 3275 | RmAb | 1:100 | MΦ |
| CD169 | Novus (Centennial, CO) | NB6005-34 | MmAb | 1:100 | MΦ, T-cell |
| PD1 | Bio SB (Santa Barbara, CA) | BSB 3151 | RmAb | 1:100 | T-cell |
| FoxP3 | Abcam (Cambridge, UK) | ab20034 | MmAb | 1:500 | Treg cell (nucleoplasm) |
| Ki-67 | Sigma-Aldrich (St. Louis, MO) | HPA 000451 | RpAb | 1:500 | Proliferation, kidney tubule; T, B-cells Nuclei |
| aSMA | Bio SB (Santa Barbara, CA) | BSB 5228 | MmAb | 1:500 | Myofiblast (cytoplasm) |
| Pan-Keratin | Abcam (Cambridge, UK) | ab234297 | RmAb | 1ug/mL | Epithelial cell (cytoplasm) |
| PTPRO | Thermo Fisher (Waltham, MA) | PAS-56964 | RpAb | 1:500 | Glomeruli (+++), (cytoplasm) |
| Kim1 | R&D Systems (Minneapolis, MN) | AF1750 | GpAb | 10ug/mL | Proximal tubule cytoplasm |
| cCas3 | Cell Signaling (Danvers, MA) | 9661-T | RpAb | 1:400 | Cytoplasmic and perinuclear |

Note: RmAb=Rabbit monoclonal antibody; MmAb=Mouse monoclonal antibody; RpAb=Rabbit polyclonal antibody; GpAb=Goat polyclonal antibody; MΦ=macrophages.

**Supplemental Table 2. Antibody Panel 2 Specifications.** Primary antibodies, clone information, dilutions, and targets for the expanded 36-marker panel deployed for high-dimensional immune phenotyping and single-cell clustering (Figure 5).

| Antibody | Vendor (City, State/Country) | Cat# | Note | Dilution factor | Location |
|---|---|---|---|---|---|
| CD1c | Thermo Fisher (Waltham, MA) | TA505411 | MmAb | 1:100 | Dendritic cells |
| CD3 | Bio SB (Santa Barbara, CA) | BSB 6452 | RmAb | 1:100 | T-cell |
| CD16 | Bio SB (Santa Barbara, CA) | BSB 3324 | RmAb | 1:100 | NK, Neutrophils, MΦ, Glomeruli |
| CD20 | Bio SB (Santa Barbara, CA) | BSB 5193 | MmAb | 1:200 | B-cell |
| CD34 | Bio SB (Santa Barbara, CA) | BSB 6488 | RmAb | 1:100 | Vascular Endothelial |
| CD36 | Thermo Fisher (Waltham, MA) | 18836-AP | RpAb | 1:1000 | MΦ, monocyte, vascular endothelium |
| CD45 | Dako (Carpinteria, CA) | M0701 | MmAb | 1:200 | Tyrosine Phosphatase, all WBC |
| CD90 | Abcam (Cambridge, UK) | ab133350 | RmAb | 1:50 | Proximal tubular & mesenchymal |
| CD163 | Bio SB (Santa Barbara, CA) | BSB 3275 | RmAb | 1:100 | MΦ |
| CD169 | Novus (Centennial, CO) | NB6005-34 | MmAb | 1:100 | MΦ |
| HIF1a | Novus (Centennial, CO) | NBP2-75978 | RmAb | 1 ug/mL | Nuclear (tubular and interstitial cells) |
| Kim1 | R&D Systems (Minneapolis, MN) | AF1750 | GpAb | 10 ug/mL | Proximal tubule cytoplasm |
| MERTK | Abcam (Cambridge, UK) | ab52968 (y323) | RmAb | 1:1000 | MΦ |
| aSMA | Bio SB (Santa Barbara, CA) | BSB 5228 | MmAb | 1:500 | Myofibroblast (cytoplasm) |
| Pan-Keratin | Abcam (Cambridge, UK) | ab234297 | RmAb | 1 ug/mL | Epithelial cell (cytoplasm) |
| PTPRO | Thermo Fisher (Waltham, MA) | PA5-56964 | RpAb | 1:500 | Glomeruli (+++), (cytoplasm) |
| cCas3 | Cell Signaling (Danvers, MA) | 9661-T | RpAb | 1:400 | Cytoplasmic and perinuclear |

Note: RmAb=Rabbit monoclonal antibody; MmAb=Mouse monoclonal antibody; RpAb=Rabbit polyclonal antibody; GpAb=Goat polyclonal antibody; MΦ=macrophages.
