## Supplementary figures and images for "Serial Immunohistochemistry for High-Dimensional Single-Cell Spatial Analysis of Human Kidney Biopsies"

### Supplemental Figure 1

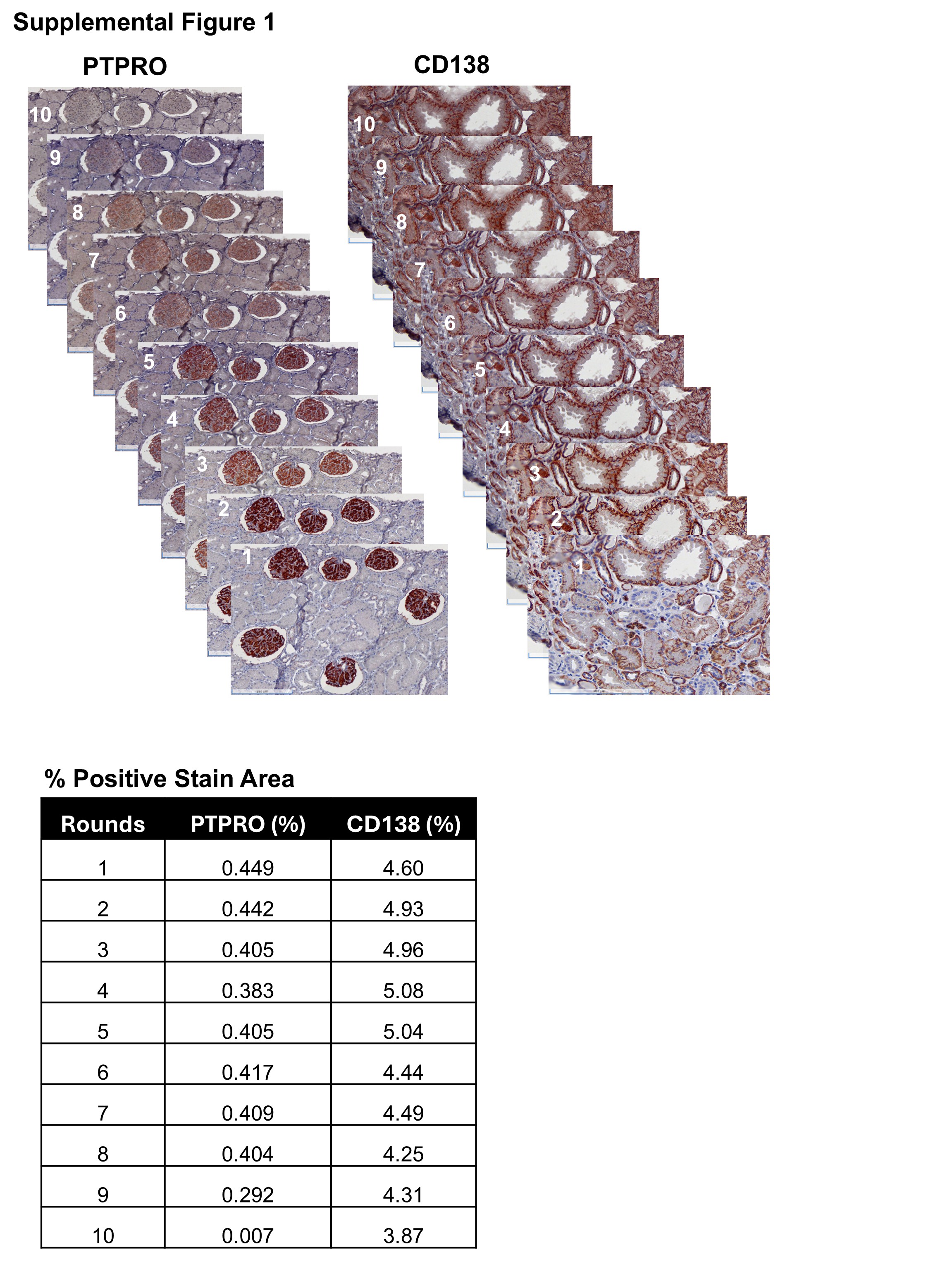
